# Assessing Oxygen Perturbation and Over-Aeration Impacts on Soluble Microbial Products (SMP) Production and Release in Activated Sludge Systems

**DOI:** 10.1101/2024.09.21.614225

**Authors:** Xueyang Zhou, Bharat Manna, Boyu Lyu, Naresh Singhal

## Abstract

Understanding the interplay between oxygen conditions and microbial activities in an activated sludge system is crucial for the optimization of wastewater treatment processes. This study explores the influence of various aeration patterns and dissolved oxygen (DO) levels on the microbial metabolic activities, with a particular focus on the synthesis and release of soluble microbial products (SMP), and the regulation of key metabolic genes and enzymes. The activated sludge system underwent different aeration patterns, including constant aeration, continuous perturbation, and intermittent perturbation under two distinct DO levels of 2mg/L and 8mg/L. We employed a combination of multi-omics techniques (metagenomics, metaproteomics, and metabolomics) along with chemical analytical methods for comprehensive sample analysis. Our results reveal an increased intracellular accumulation of amino acids and enhanced release of protein under the conditions of oxygen perturbation. Furthermore, elevated DO levels fostered the accumulation of poly-3-hydroxybutyrate intracellularly and the release of protein and fatty acids as SMP. This outcome is associated with the abundance of key metabolic genes and enzymes, thereby highlighting the metabolic flexibility of microbes under different oxygen conditions. These findings offer valuable insights into microbial metabolic dynamics under varying oxygen conditions, thereby providing guidance for more efficient and sustainable strategies in wastewater treatment and resource recovery.

**SYNOPSIS:** This study explores the complex interplay between varying oxygen conditions and microbial activity in activated sludge systems, focusing on the impact on the synthesis and release of soluble microbial products (SMP). Findings highlight the significant influence of oxygen perturbation and dissolved oxygen levels on protein synthesis and fatty acid metabolic processes.

## 1. INTRODUCTION

In traditional wastewater treatment paradigms, the primary goal has been the removal and disposal of pollutants from wastewater to safeguard public health and protect water bodies from contamination. However, this linear approach is increasingly seen as unsustainable due to the high energy consumption, valuable resources lost, and greenhouse gas emissions in the process.^1,2^ As a consequence, the field is shifting towards a more circular and sustainable approach - considering wastewater not as a waste but as a resource.^3–6^ This new paradigm aims to recover valuable components such as safe water, energy, nutrients, and biopolymers, thereby contributing to the sustainable development of human societies and the environment.^7,8^

The activated sludge process, one of the most widely used biological treatment methods, provides an ideal platform for resource recovery from wastewater.^9,10^ In this process, diverse microorganisms form flocculent aggregates, known as activated sludge, that metabolize organic and inorganic pollutants, transforming them into microbial biomass and carbon dioxide.^11^ Beyond this pollutant removal function, activated sludge microorganisms can also synthesize valuable intracellular storage compounds, such as amino acids, fatty acids, and polyhydroxyalkanoates (PHA), and release extracellular organic matter like soluble microbial products (SMP) which can be harnessed as resources.^12,13^ Given this potential, it is essential to understand and optimize the operational conditions of the activated sludge process to improve resource recovery efficiency.

A key operational factor in the activated sludge process is the aeration condition, which influences the metabolism of the microbial community and thereby affects the production and release of intracellular and extracellular compounds.^14^ Current understanding of how varying aeration conditions, like oxygen perturbation and over-aeration, might influence the specific metabolite production, accumulation, and release mechanisms within the activated sludge process is still limited.^15^ These gaps in knowledge hinder our ability to strategically manipulate aeration conditions to optimize resource recovery from wastewater.

This study investigated the effect of different oxygen conditions on the microbial release of SMP in an activated sludge system, focusing on the synthesis and release of specific metabolites and the role of key metabolic genes and enzymes. The activated sludge system was exposed to diverse aeration patterns encompassing constant aeration (CA), continuous perturbation (CP), and intermittent perturbation (IP), under two distinct dissolved oxygen (DO) levels of 2mg/L and 8mg/L. The samples were meticulously analyzed employing a combination of chemical analytical techniques (nitrogen concentration determination, organic matter fluorescence spectroscopy) and multi-omics analysis approaches (metagenomics, metaproteomics, and metabolomics).

Our findings revealed that changes in aeration conditions affected the intracellular accumulation of amino acids and fatty acids, and extracellular release of proteins and fatty acids, as well as the abundance of key metabolic genes involved in PHA synthesis and degradation. These results suggest that strategic manipulation of aeration conditions could promote the accumulation and release of valuable compounds from activated sludge, contributing to resource recovery from wastewater.

## 2. MATERIALS AND METHODS

### 2.1 Experimental Setup and Operation

The activated sludge for this study was taken from the Mangere municipal wastewater treatment plant in Auckland, New Zealand. The 1 L initial reaction system included activated sludge (3g/L MLSS), inorganic carbon (3840 mg NaHCO_3_), and 1 mL of trace element solution (Table S1). The reaction was processed at a room temperature of 20±1 for 48 hours. The biosystem was mixed by a magnetic stirrer and exposed to different aeration conditions by an automatic DO control system. Six aeration conditions include three different modes and two different levels. Specifically: Continuous Perturbation (0.1 to 2.0 mg/L or 0.1 to 8.0 mg/L), Intermittent Perturbation (0 to 2.0 mg/L or 0 to 8.0 mg/L, aerobic time: anoxic time = 1:1) and Continuous Aeration (2 mg/L or 8 mg/L).

After the reaction began, the concentrated artificial wastewater (Table S2) containing organic carbon source (methanol), inorganic nitrogen source (ammonia nitrogen), and phosphorus source was fed into the reaction system at an even rate. Depending on the rate of nutrient consumption by the biosystem, a total of 1600 mg COD, 320 mg NH_4_ ^+^-N for IP or 400 mg NH_4_ ^+^-N for CP and CA, and 40 mg PO ^3-^-P was continuously injected by a syringe pump for 48 hours.

Originally, we planned to conduct three biological replicates for each experimental condition. However, due to unforeseen circumstances related to the sudden closure of the laboratory in response to the COVID-19 pandemic, we regretfully could not complete the 48-hour sampling under the DO8 conditions. Despite this setback, we ensured that three biological replicates were maintained for all other time points.

### 2.2 Carbon and Nitrogen Analysis Methods

The water samples taken in the experiment were filtered and analysed for water quality, NH_4_ ^+^- N and COD were determined by commercial reagent vials (Hach, USA). Total organic carbon (TOC), total inorganic carbon (TIC) and total nitrogen (TN) were measured by a TOC analyser (Shimadzu, Japan). NO_2_ ^-^-N and NO_3_ ^-^-N were determined by ion chromatography (Thermo Fisher, USA).

The amount of dissolved organic nitrogen was determined by the difference between TN and total inorganic nitrogen (NO_2_^−^-N, NO_3_^−^-N and NH_4_^+^-N). The gaseous nitrogen production was obtained by subtracting the dissolved and biomass nitrogen from NH_4_ ^+^-N consumed by the biosystem. Biomass nitrogen was obtained from MLSS data by the accepted empirical formula for bacterial cell (C_5_H_7_O_2_N).^16^ MLSS was analyzed according to the standard methods.^17^

### 2.3 SMP Measurement

The three-dimensional excitation-emission matrix (3D-EEM) analysis method was used for the determination of fluorescent organic substances in the samples by an Aqualog A-TEEM Spectrometer (Horiba, Japan). Excitation wavelengths were set from 240 to 600 nm at interval of 3 nm; emission wavelengths were set from 212.70 to 622.21 nm at interval of 3.28 nm. The fluorescence from 3D-EEM was modelled using PARAFAC by stardom.^18^

The extracellular protein concentration was concentrated 40-fold by freeze-drying before being measured by the RC DC™ Protein Assay Kit I (Bio-Rad, USA). The standard curve was established using the bovine γ-globulin standard.

### 2.4 Metabolomics

The intracellular metabolites’ relative abundance in activated sludge was quantified using a GC/MS platform. For all conditions, three biological replicates were taken from the 24-hour samples. At the 48-hour mark, three biological replicates were acquired for the DO2 conditions (CA2, CP2, and IP2), and two for the DO8 conditions. Each sample with two technical replicates. Sludge samples (10 mL) were centrifuged, quickly quenched in liquid nitrogen, and preserved at -80 °C for subsequent processing. Sludge pellets were mixed with a cold methanol-water solution and an internal standard, 2,3,3,3-d4-alanine. Metabolites were released via repeated freeze-thawing and rigorous shaking, with the supernatant collected via centrifugation. Another round of metabolite extraction occurred with more methanol-water solution. Resulting metabolite extracts were freeze-dried and derivatized using methyl chloroformate (MCF) and fatty acid direct transesterification (FAMEs).

For the analysis of amino acids, MCF derivatization was used. The metabolite extract was resuspended in 400 µL of 1 M sodium hydroxide solution, and 68 µL of pyridine and 334 µL of methanol were added. MCF (40 µL) was added twice, with each addition followed by 30 seconds of vigorous agitation. The mixture was then combined with 400 µL of chloroform and agitated for 10 seconds. Afterward, 800 µL of a 50 mM bicarbonate solution was added, and the mixture was stirred and centrifuged. The aqueous phase was discarded, and any trace of water in the chloroform phase was removed using anhydrous sodium sulfate.

The MCF derivatives were analyzed using an Agilent GC7890 GC-MS system coupled with an MSD597 unit. This system was equipped with a ZB-1701 GC capillary column (30 m × 250 µm (id) × 0.15 µm film thickness) and a 5 m guard column (Phenomenex, Torrance, CA, USA). The generated GC-MS chromatograms were deconvoluted by AMDIS software (NIST, Boulder, CO, USA) and compared against the in-house MCF MS library to identify metabolites.^19^

For the analysis of fatty acids, FAMEs derivatization was used. The metabolite extract was placed in a sealable borosilicate tube, to which nonadecanoic acid (4µg/L) was added as an internal standard. This was followed by the slow addition of 200 µL of acetyl chloride. After sealing the tube, it was subjected to a reaction at 100°C for one hour. After cooling, a 5 mL K2CO3 solution (6%) was added to neutralize the mixture. The tube was then shaken, centrifuged, and a portion of the toluene upper phase was extracted for GC-MS analysis and quantification using a fatty acid standard curve.^20^

Metabolite data filtering was facilitated by the “MassOmics” R package, relying on MS spectra and retention times of the derivatized metabolites.^21^ Metabolite abundance normalization was performed using d4-Alanine, and baseline calibration was achieved by subtracting blank information from metabolite abundance in samples. The relative abundance of intracellular metabolites was eventually derived by analyzing the difference between initial and 48-hour sample data. Data visualization was completed using MetaboAnalyst 5.0,^22^ and ComplexHeatmap.^23^

### 2.5 Metaproteomics

This study entailed metaproteomic analysis on sludge samples across distinct experimental conditions, employing three biological replicates were taken at the 24-hour mark. At the 48-hour point, three biological replicates were gathered for DO2 conditions, while for DO8 conditions, two biological replicate was obtained. Sludge samples were subjected to pelleting, washing, and centrifugation at 6,113 x g for 5 minutes at 4 °C. Resulting pellets were instantaneously frozen in liquid nitrogen and preserved at -80 °C. Proteins were extracted by resuspending the pellet in lysis buffer (pH 8) and sonication. The resulting lysate was mixed with 20% trichloroacetic acid in acetone, vortexed, incubated, and centrifuged. After discarding the supernatant, the pellet was subjected to consecutive washing steps with acetone and 80% acetone in water. The protein pellet was air-dried and resuspended in 50 mM Tris buffer.

Protein purification and quantification were conducted using SpeedBead Carboxylate-Modified E3 and E7 magnetic particles (Sera-Mag, USA) and the EZQ protein assay kit (Invitrogen, USA), respectively. The sample underwent reduction and alkylation via DTT and iodoacetamide, respectively. Trypsin was added to perform protein digestion. Post-digestion, the sample was diluted and filtered using a Vivaspin centrifugal concentrator. Subsequent extraction of peptides passed through was performed with Oasis Prime HLB 1cc (30 mg) solid-phase extraction. The extracted peptides were concentrated and prepared for nano LC-MS/MS analysis.

The samples were desalted and segregated using a NanoLC 400 UPLC system (Eksigent, USA). Mass spectrometry analysis was accomplished via the TripleTOF 6600 Quadrupole-Time-of-Flight mass spectrometer (Sciex, USA). Lastly, the data was compared using Protein Pilot version 5.0 (Sciex, USA) against the database established by metagenomics results of the samples.

### 2.6 Metagenomics

For DNA extraction, 10 mL sludge samples were collected. Three biological replicates were obtained from 24-hour samples across all conditions. Additionally, three biological replicates were obtained from the 48-hour samples under DO2 conditions. However, under the DO8 conditions at the 48-hour mark, only a single biological replicate was collected. The extraction process was executed according to the protocol of DNeasy PowerSoil Kit (Qiagen, German), and the extracted DNA was stored at -20°C for subsequent analysis.

The extracted DNA samples were then dispatched to the Auckland Genomics Centre for metagenomic analyses. The laboratory procedures included concentrating the DNA samples, performing a quality control (QC) check, and preparing the DNA for Illumina sequencing, which included polymerase chain reaction (PCR) and indexing. Following the library QC, normalization was performed, and a final pool check was conducted using a bioanalyzer. Finally, sequencing was performed on a HiSeq platform, generating approximately 400 million 2x150bp paired-end reads. The data returned from the sequencing process were analyzed using SqueezeMeta.^24^

## 3. RESULTS

### 3.1 Oxygen Perturbations and Over-aeration Promote SMP Release

#### 3.1.1 The Dissolved Organic Nitrogen in Extracellular Water Samples

An activated sludge system was continuously operated under various aeration conditions over a period of 48 hours. Different nitrogen-related parameters (total nitrogen, ammonia nitrogen, nitrite nitrogen, nitrate nitrogen) were measured in the water samples to determine the quantities of ammonia nitrogen that had been transformed into various forms of nitrogen (gaseous nitrogen, nitrite, nitrate, and organic nitrogen) (Figure S2).

The concentration of organic nitrogen in the extracellular water samples was derived by calculating the difference between the total nitrogen and total inorganic nitrogen (Figure 1a). Comparison of data from different samples revealed that the secretion efficiency of dissolved organic nitrogen (DON) synthesized in the activated sludge system was contingent upon the aeration conditions.

**Figure 1.**
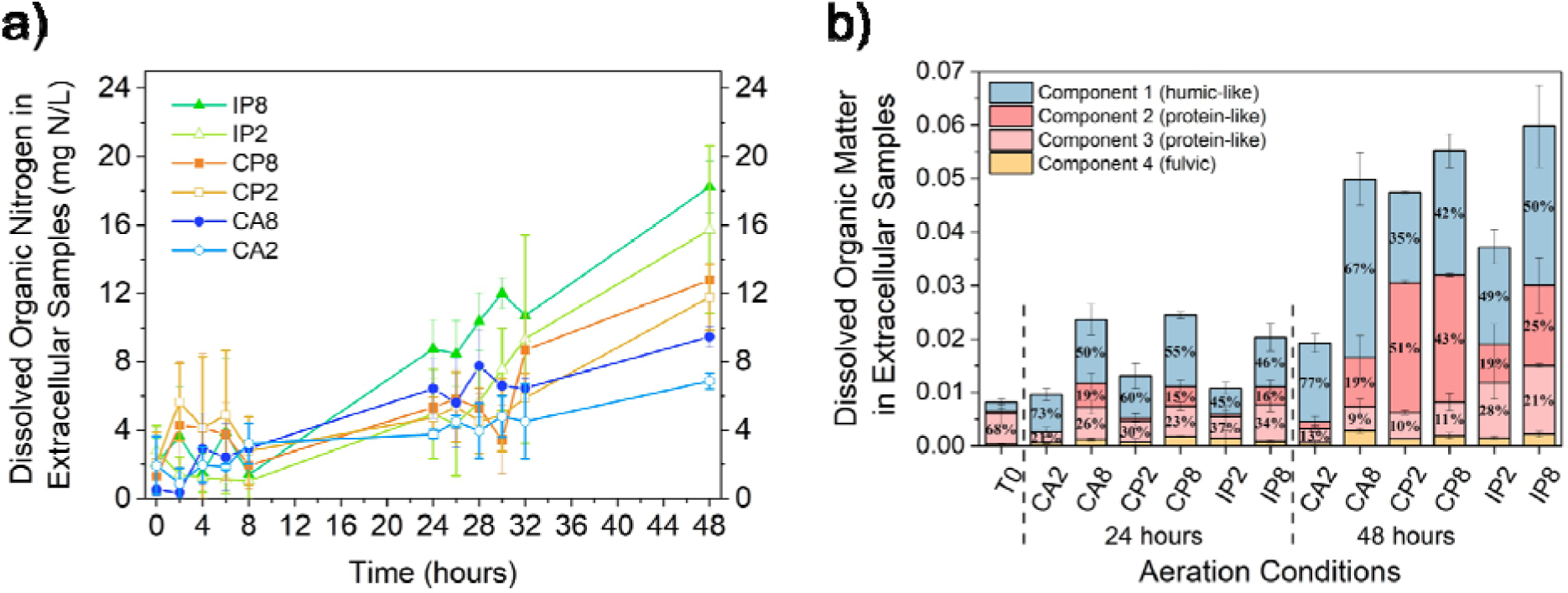
Different aeration parameters affected dissolved organic nitrogen in extracellular samples of activated sludge systems: (a) The dissolved organic nitrogen in extracellular samples of activated sludge system under different aeration conditions; (b) The relative concentration of fDOM components in extracellular samples. Error bars represent standard deviations (CA2, CP2, and IP2 have biological triplicates; n=3. CA8, CP8, and IP8 have biological duplicates; n=2).

Among the three different aeration patterns, the oxygen perturbation conditions (IPs and CPs) led to higher DON concentrations in extracellular water samples than the CAs. Specifically, under IP2 and IP8 conditions, the activated sludge system produced DON concentrations of 15.75±4.91 mg N/L and 18.24±1.50 mg N/L, respectively, in the extracellular water samples after 48 hours of operation. Under CP2 and CP8, they were 11.78±1.94 mg N/L and 12.79±0.02 mg N/L. Moreover, the data suggest that elevated DO level can enhance the release of more organic matter from the activated sludge. This trend was consistently observed across all aeration patterns, including CA, CP, and IP.

#### 3.1.2 Composition and Content of DOM in Extracellular Water Samples

In order to assess the dissolved organic matter (DOM) produced and extruded by microbes, fluorescence-based organic substances were identified in the extracellular water samples from the activated sludge system via 3D-EEM. The differentiation in component types within samples is chiefly manifested by disparities in the locational shifts of fluorescence peaks within the 3D-EEM spectra.^25^ Figure S3 illustrates the spectral peak localities that correlate to four organic matter constituents present in the extracellular water samples. The content of each component within the samples can be estimated from the fluorescence intensities at various peak positions using the PARAFAC model. Figure 1b demonstrates the diverse constituents and their corresponding quantities in the organic matter from the extracellular water samples, acquired from the activated sludge system under differing aeration scenarios at three distinct time points. It is evident that the volume of organic matter within the extracellular environment progressively amplifies over time.

Of all the aeration conditions, the activated sludge system under CA2 exhibited the minimum quantity of DOM release into the water body. In contrast, the microbial systems operating under oxygen perturbation showcased elevated extracellular DOM content post 48 hours of operation. Moreover, irrespective of the 24-hour or 48-hour operational duration, over-aerated samples, consistently manifested heightened DOM content. This observation corroborates the previously deduced data regarding extracellular organic nitrogen concentrations.

Furthermore, the predominant constituents of extracellular DOM exhibited noticeable discrepancies across the various aeration patterns. Under the CA conditions, C1 emerged as the primary component, reaching 77% and 67% in CA2 and CA8, respectively. Yet, under the CP conditions, the proportion of C2 attained its peak, comprising 51% under CP2 and 43% under CP8. In the extracellular water samples under the IP conditions, while C1 remained the most abundant, intriguingly, the proportion of C3 exceeded that in other conditions. Such differences in DOM component proportions suggest that different aeration patterns might impact the compositional variation of SMP excreted by the activated sludge system, possibly attributable to the variable metabolic activity of microbes. Alterations in the DO concentration might induce variable metabolic conditions for diverse bacterial species within the system, thereby modulating the metabolic pathways associated with SMP generation and excretion.

### 3.2 Influence of Oxygen on Amino Acid Metabolism

#### 3.2.1 Extracellular Protein and Intracellular Amino Acids

Protein-like compounds, encompassing proteins, amino acids, oligopeptides, enzymes, and other nitrogen-containing compounds, are valuable bioproducts in biomass resource recovery processes.^26–30^ The extent to which activated sludge releases proteins into the water under varying aeration conditions can be assessed through analysis of protein concentrations in extracellular water samples taken at diverse points throughout operation. Figure 2a presents the protein concentrations in the water samples over a 48-hour operational period under six distinct aeration conditions. Notably, as the reactor operation proceeds, the activated sludge system under oxygen perturbation conditions tends to yield more extracellular proteins, with the effect being more pronounced under CP compared to IP. Furthermore, elevated DO conditions can foster increased protein release from the cells.

**Figure 2.**
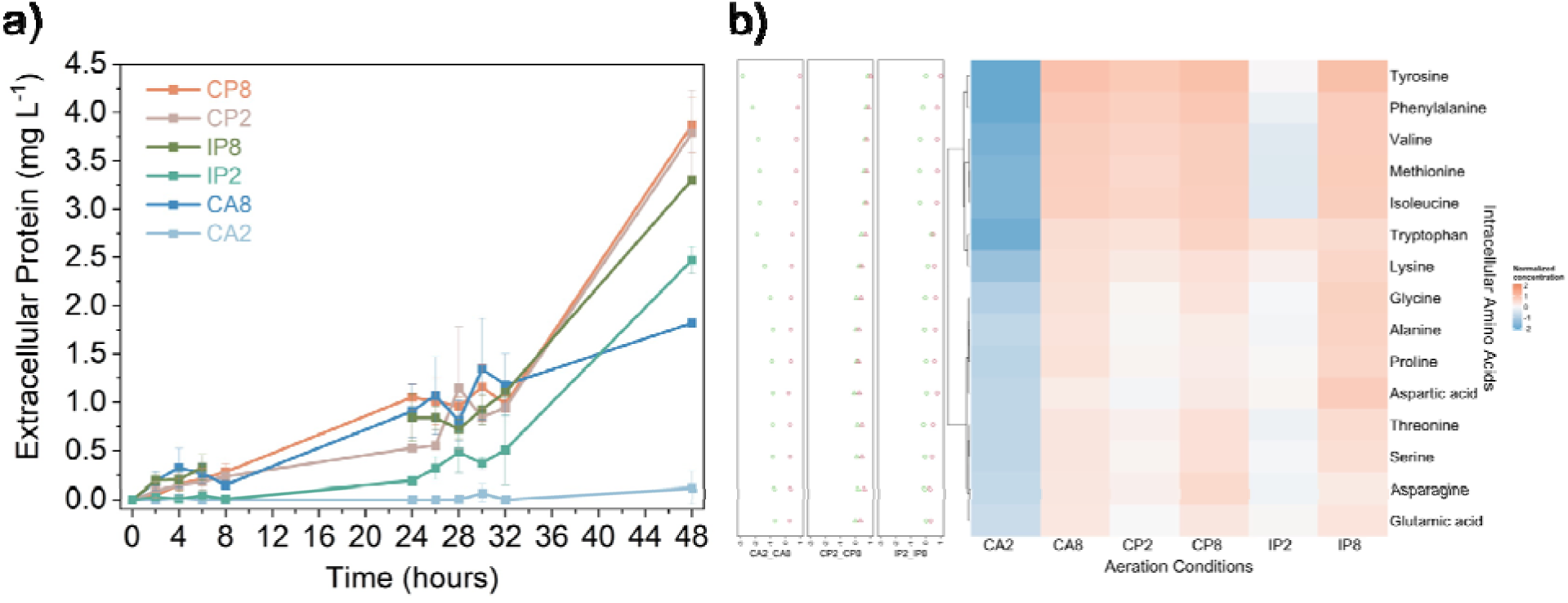
Oxygen perturbation and over-aeration promoted extracellular protein release and intracellular amino acid abundance: (a) The concentration of protein in the extracellular water samples. Error bars represent standard deviations (CA2, CP2, and IP2 have biological triplicates; n=3. CA8, CP8, and IP8 have biological duplicates; n=2); (b) The abundance of intercellular amino acids in the activated sludge system exposed to different aeration strategies for 48 hours.

Amino acids serve as fundamental building blocks in the synthesis of protein-like compounds.^31^ To determine the influence of aeration conditions on amino acid metabolism, an MCF-derivatization metabolomics approach was employed to quantify amino acid abundance within biomass samples. Figure 2b displays the process of synthesizing and accumulating amino acids intracellularly from a single organic carbon source (methanol) and inorganic nitrogen (ammonia), under six different aeration conditions over 48 hours. Among the three conditions with the DO2 level, the oxygen perturbation conditions (CP2 and IP2) induced more intracellular amino acid accumulation than CA2. More strikingly, under over-aeration conditions, irrespective of the aeration pattern, the abundance of common amino acids within the biomass exceeded that under standard aeration conditions.

To investigate whether variations in the proportions of different protein-like compound components within the extracellular DOM under different aeration patterns relate to the status of intracellular amino acid accumulation, the trends in intracellular amino acid abundance were analyzed for each aeration pattern. Beginning with the representative CP conditions, the abundance trends of intracellular amino acids under CP2 and CP8 were largely parallel, aligning with the near-identical DOM component composition observed in the 3D-EEM analysis.

In the case of IP conditions, the abundance of certain intracellular amino acids remained consistent across both conditions, although some displayed higher abundance under IP8. In harmony with the 3D-EEM analysis outcomes, the DOM under both IP conditions harbored a higher fraction of the C3 component, but the C2 component proportion was elevated in IP8 compared to IP2.

For the CA conditions, considerable disparities were observed in intracellular amino acid abundance, with all types of amino acids manifesting obviously higher abundance under CA8 compared to CA2. Similarly, CA8 also displayed a higher proportion of protein-like compound components within the DOM than CA2.

The data is normalized by internal standard, Log10 transformation, and Pareto scaling. The heatmap and dot plots show the average value of the same condition. Row clustering according to ‘Ward’. In the dot plots, green dots represent CA2/CP2/IP2 and red dots represent CA8/CP8/IP8 (CA2, CP2, and IP2 have technical duplication results for biological triplicates; n=6. CA8, CP8, and IP8 have technical duplication results for biological duplicates; n=4).

#### 3.2.2 The Abundance of Aminoacyl-tRNA Synthetases

Aminoacyl-tRNA synthetases (aaRS) play a pivotal role in the protein synthesis process by associating specific amino acids with their respective transfer RNAs (tRNAs) - a key step in translating genetic information into functional proteins.^32^ Given that each aaRS is tailor-made for a unique amino acid and tRNA pairing, the relative abundance of a particular aaRS may well signify the frequency of its corresponding amino acid usage in microbial protein synthesis. Consequently, fluctuations in the abundance of different aaRS could hint at the changing requirements for various proteins by microbes under assorted environmental conditions.

With an aim to explore whether the higher percentage of the C3 component in protein-like substances released by microbial systems under intermittent aeration (IP conditions), as compared to constant aeration (CA) and continuous positive aeration (CP conditions), an indication of an adaptive adjustment in protein synthesis strategies, we conducted a metaproteomics analysis of activated sludge samples. These samples were treated under different aeration conditions for a duration of 48 hours. We observed intriguing variations in the abundance of various aaRS across the different samples (Figure 3), with certain types of aaRS being notably dominant under IP conditions - a characteristic that was different from those under CA and CP conditions. This observation suggests that the intermittent periods of anoxia in the aeration regime might have nudged the activated sludge system to modify protein synthesis strategies, leading to the release of distinct protein-like compounds.

**Figure 3.**
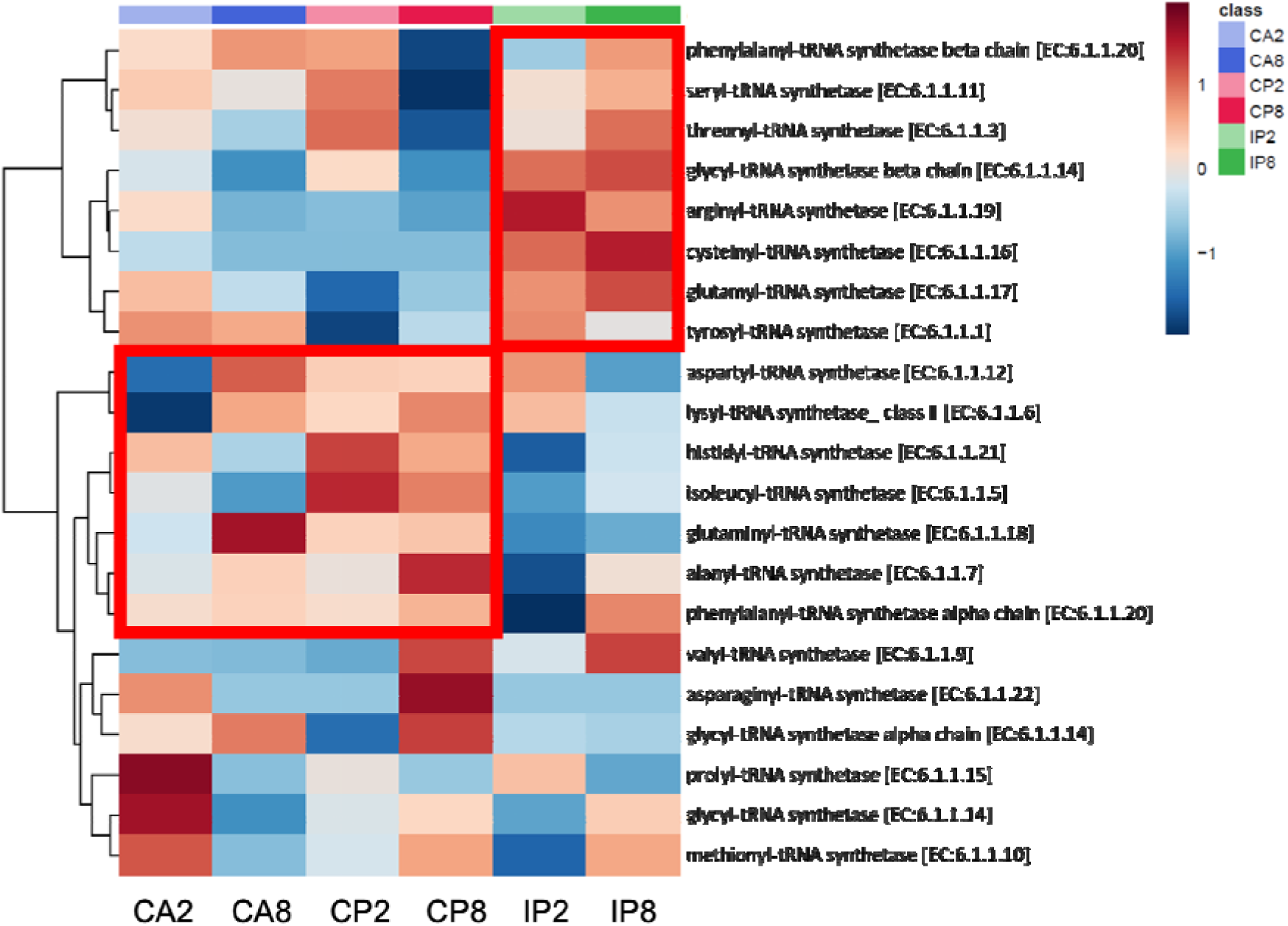
The abundance of various aminoacyl-tRNA synthetases under different aeration conditions. The data is normalized by Log10 transformation and Pareto scaling. The heatmap shows the average value of the same condition. Row clustering according to ‘Ward’ (CA2, CP2, and IP2 have biological triplicates; n=3. CA8, CP8, and IP8 have biological duplicates; n=2).

### 3.3 Oxygen Influence on Fatty Acid Metabolism

#### 3.3.1 Extracellular and Intracellular Fatty Acids

The humic-like substances present in SMP denote a category of organic compounds possessing qualities and functionalities reminiscent of humic matter, typically residing in natural environments such as soils, peat, aquatic ecosystems, among others.^33^ These substances serve a pivotal agents in soil amelioration and environmental safeguarding, enhancing plant nutrient uptake, stabilizing soil and water pH, and adsorbing deleterious substances.^34–36^ Fatty acids, within this realm, play an indispensable role.^37^ To examine the production and release mechanisms of extracellular fatty acids in greater depth, we employed a metabolomics approach grounded in FAMEs derivatization, facilitating the quantification of fatty acids in both the extracellular aqueous and biomass specimens of the activated sludge system.

Figure 4 illustrates the relative abundance of 14 extracellular and 19 intracellular fatty acids. Predominantly, the detected fatty acids were long-chain fatty acids (LCFA), characterized by carbon chain lengths spanning 13 to 21 carbon atoms. Of these, 9 were detected extracellularly, and 14 were intracellular. Moreover, out of the 12 fatty acids detected in both intra- and extracellular environments, 11 were LCFA. This evidence underscores that whether accumulated internally or released externally, the fatty acids are indeed predominantly LCFA.

**Figure 4.**
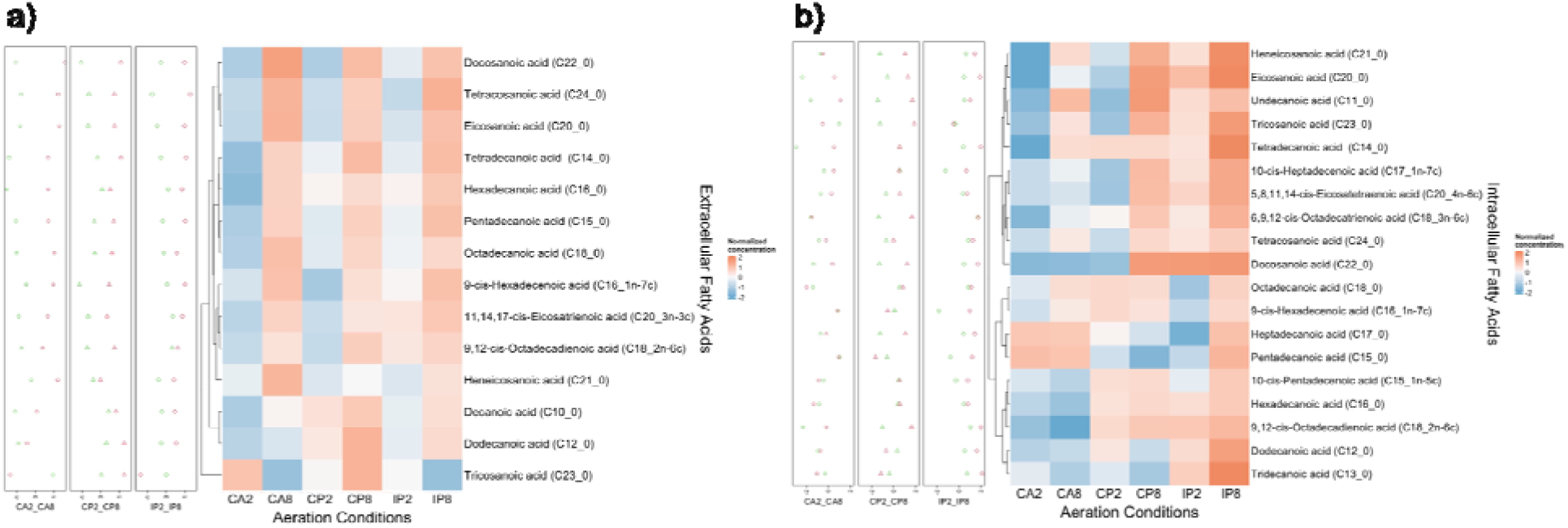
Over-aeration promoted the release of extracellular fatty acids and the accumulation of intracellular fatty acids: (a) and the biomass samples (b) of the activated sludge system exposed to different aeration strategies for 48 hours. The data is normalized by internal standard, Log10 transformation, and Pareto scaling. The heatmap and dot plots show the average value of the same condition. Row clustering according to ‘Ward’. In the dot plots, green dots represent CA2/CP2/IP2 and red dots represent CA8/CP8/IP8 (CA2, CP2, and IP2 have technical duplication results for biological triplicates; n=6. CA8, CP8, and IP8 have technical duplication results for biological duplicates; n=4).

Upon comparing the abundance of extra- and intracellular fatty acids under disparate aeration conditions, a shared pattern emerged; over-aeration consistently led to a heightened abundance of fatty acids. This trend could be attributed to variations in fatty acid synthesis and utilization efficiency under high versus low DO conditions. Unlike the scenario involving amino acids, the abundance of extracellular or intracellular fatty acids did not show clear differences under the three DO2 aeration conditions. Notably, this observation aligns with their similar proportions of humic-like components within the DOM fraction.

The results of metabolomics analysis, detailing the abundance of amino acids and fatty acids, corroborate our experimental observations derived from the 3D-EEM spectrophotometry. Specifically, oxygen disturbances foster the accumulation of intracellular amino acids and protein release, while CP and IP induce differential protein-like fractions. Furthermore, over-aeration can simultaneously augment the accumulation of intracellular amino acids and fatty acids, along with the liberation of protein-like and humic-like substances.

#### 3.3.2 The Fatty Acid Biosynthesis Metabolism

To gain a more in-depth understanding of the microbial regulatory mechanisms connected with the intra- and extracellular dynamics of fatty acids, we conducted a comprehensive examination of the abundance of relevant genes and enzymes implicated in fatty acid synthesis and bio-utilization. This was achieved through a dual approach using metagenomics and metaproteomics.

Within the metabolic pathways of fatty acid biosynthesis, Acetyl-CoA carboxylase (EC: 6.4.1.2) functions as a critical enzyme. It facilitates the conversion of Acetyl-CoA and CO_2_ into Malonyl-CoA, a pivotal initiating factor for fatty acid synthesis. Six homologous genes, specifically accA, accB, accC, accD, bccA, and accD6, are associated with this enzyme.^38,39^

During the 24th and 48th operational hours, the activated sludge samples showed that, of these six genes, accA, accB, accC, and accD had a higher abundance compared to accD6 and bccA (Figure S12, S13). This may be due to the greater functional importance of these four genes in encoding the ACC enzyme complex. Among the three patterns, IP appears to have a slightly lower gene abundance, though the difference is not substantial.

Upon specific evaluation of the Acetyl-CoA carboxylase enzyme abundance in the activated sludge system (Figure 5a), irrespective of the DO level, systems employing the CP consistently displayed a higher abundance of the ACC enzyme complex, followed by CA, and then IP. This indicates that activated sludge systems subjected to continuous and rapid oxygen fluctuations can induce an elevated ACC enzyme abundance, thereby fostering fatty acid synthesis from the perspective of enzymatic reaction efficacy. In contrast, the IP approach exhibits reduced enzymatic efficiency, likely due to ATP scarcity resulting from an insufficient oxygen supply.

**Figure 5.**
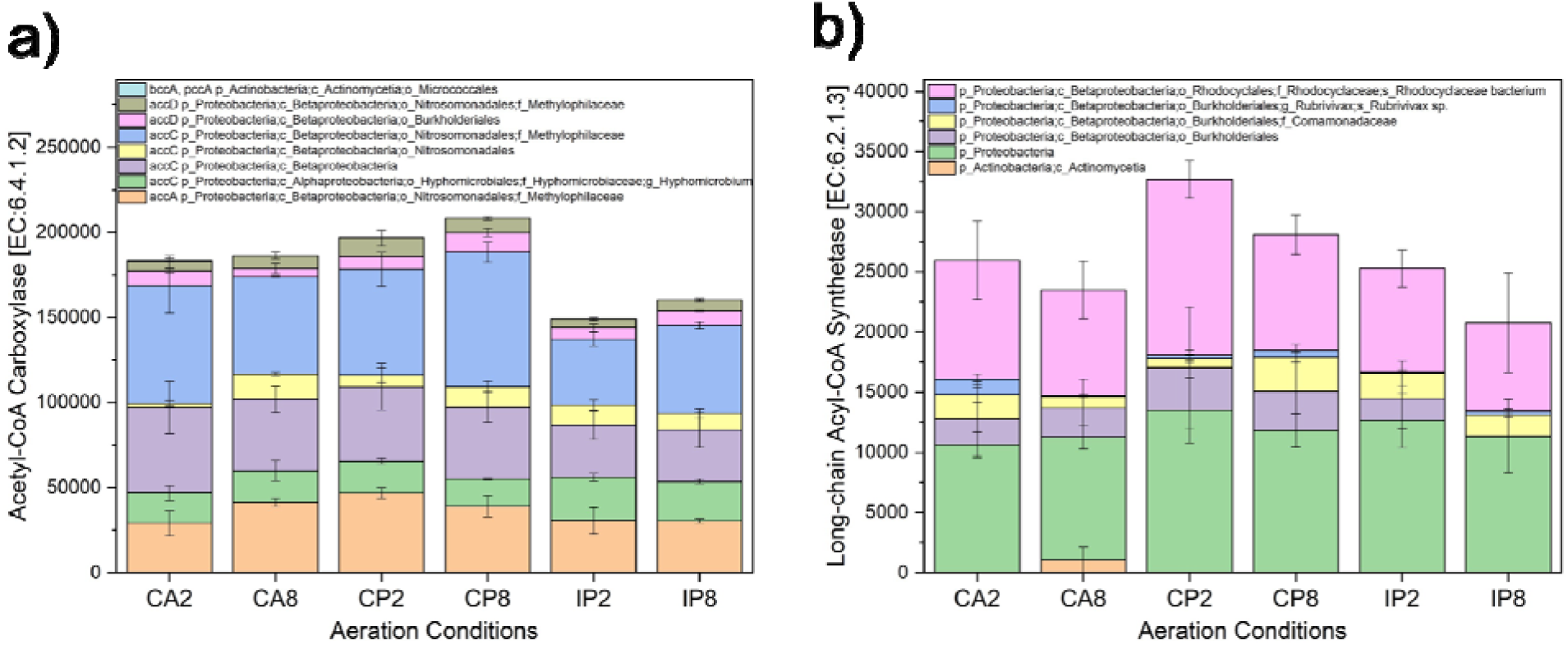
Key enzymes for fatty acid production and utilization contribute to the relative abundance of fatty acids: (a) Abundance variations of Acetyl-CoA carboxylase under six aeration conditions after 48 hours of operation. The error bars represent standard deviations; (b) Abundance variations of long-chain Acyl-CoA synthetase under six aeration conditions after 48 hours of operation. The error bars represent standard deviations (CA2, CP2, and IP2 have biological triplicates; n=3. CA8, CP8, and IP8 have biological duplicates; n=2).

For each of the three aeration patterns, higher DO levels corresponded to a marginally increased ACC enzyme abundance. This suggests that enhancing the oxygen concentration during steady aeration or broadening the oxygen concentration range during oxygen fluctuations can both stimulate fatty acid synthesis. This finding is in agreement with the relative abundance trend of intracellular fatty acids identified via our metabolomics analysis.

#### 3.3.3 The Fatty Acid Utilization Metabolism

Adopting the same analytical approach as before, we directed our focus towards the key node in the metabolic pathway of fatty acid utilization. The initiation of fatty acid utilization in biological systems hinges on their activation into CoA derivatives, thus enabling their engagement in metabolic pathways such as α-and β-oxidation, elongation, and phospholipids formation.^40–42^ This pivotal activation step is conducted by the acyl-CoA synthetase (ACS) family. Given that the length of the fatty acid carbon chain determines its hydrophobicity and solubility, there are different ACS enzymes within the ACS family, each specialized for the activation of fatty acids of varying chain lengths.^43^

We undertook an analysis of metagenomic data corresponding to distinct ACS members within the ACS family, each possessing differential activation preferences. Nine distinct members of the ACS family were identified and analyzed for their relative abundance, each of these members corresponds to a distinct gene within the ACS family (Table S5).^44^ More microorganisms demonstrate the presence of genes K01895, K01908, and K01897 (Figure S14, S15). This suggests that, in terms of growth, genes facilitating the utilization of short-chain and long-chain fatty acids are more commonly present in the microbial communities within the activated sludge systems, as opposed to other genes related to fatty acid utilization.

Analyzing from an enzymatic regulatory perspective, over-aeration appeared to decrease the abundance of Acyl-CoA synthetase long-chain family member (ACSL) (Figure 5b). Interpreting these observations in conjunction with the prevalence of LCFA, it appears that under over-aeration conditions, enhanced oxygen uptake may have facilitated a more optimal ATP production environment. This likely prompted the microorganisms to boost the efficiency of fatty acid synthesis by upregulating the expression of ACC enzyme. However, these conditions seemed to simultaneously inhibit the expression of ACSL, thus decreasing the utilization efficiency of long-chain fatty acids. This phenomenon could potentially contribute to an increased accumulation of long-chain fatty acids within the cells.

LCFA may also derive from carbon chain shortening in very long-chain fatty acids (VLCFA). The gene K08746 is responsible for the utilization of VLCFA, which belongs to the ACS subfamily, demonstrates a sensitivity to high oxygen conditions (Figure S14, S15). The biological utilization of VLCFA is considerably hampered under high DO levels, potentially leading to an accumulation of VLCFA. To detoxify excess VLCFA, they are subjected to peroxisomal β-oxidation, which shortens the carbon chain.^45^ This process relies critically on a bifunctional protein that possesses both enoyl-CoA hydratase and hydroxyacyl-CoA dehydrogenase activities.^46^ High DO conditions trigger an enhanced expression level of this enzyme across all aeration patterns tested in the six different aeration conditions (Figure S16). The highest expression levels, observed under IP8 and CA8 conditions, correspond with a higher abundance of intracellular LCFA and VLCFA, as indicated by metabolomics results. These findings suggest that high DO aeration conditions, whether under perturbed or non-perturbed oxygen circumstances, could interfere with the biological utilization of VLCFA, contributing to their intracellular accumulation. During the microbial detoxification process, LCFA and VLCFA may be released from the cell, leading to an elevated concentration in the water.^47,48^

### 3.4 Over-aeration Promotes PHA Accumulation

#### 3.4.1 The PHA Content in Biomass Samples

Polyhydroxyalkanoates (PHA) are biodegradable polymers synthesized by microorganisms. Typically, bacteria induce and accumulate PHA in response to nutritional deficiencies and environmental stressors by modulating the gene expression of the PHA synthesis metabolic pathway.^49,50^ Poly-3-hydroxybutyrate (PHB), a short-chain PHA composed of 3-hydroxybutyrate monomers, is the most prevalent type of PHA in many bacteria.^51,52^ By quantifying the PHB content in activated sludge biomass samples under various aeration conditions, we found that oxygen perturbations and over-aeration led to increased PHB levels (Figure 6a, 6b). Under CP8 and IP8 conditions, the PHB content reached 11.82±0.17% and 12.52±0.23%, respectively, at the 24-hour mark and 12.95±0.30% and 12.42±2.28% after 48 hours of operation. In contrast, the CA2 condition resulted in PHB contents of 7.77±0.27% and 9.14±0.46% at 24 and 48 hours, respectively.

**Figure 6.**
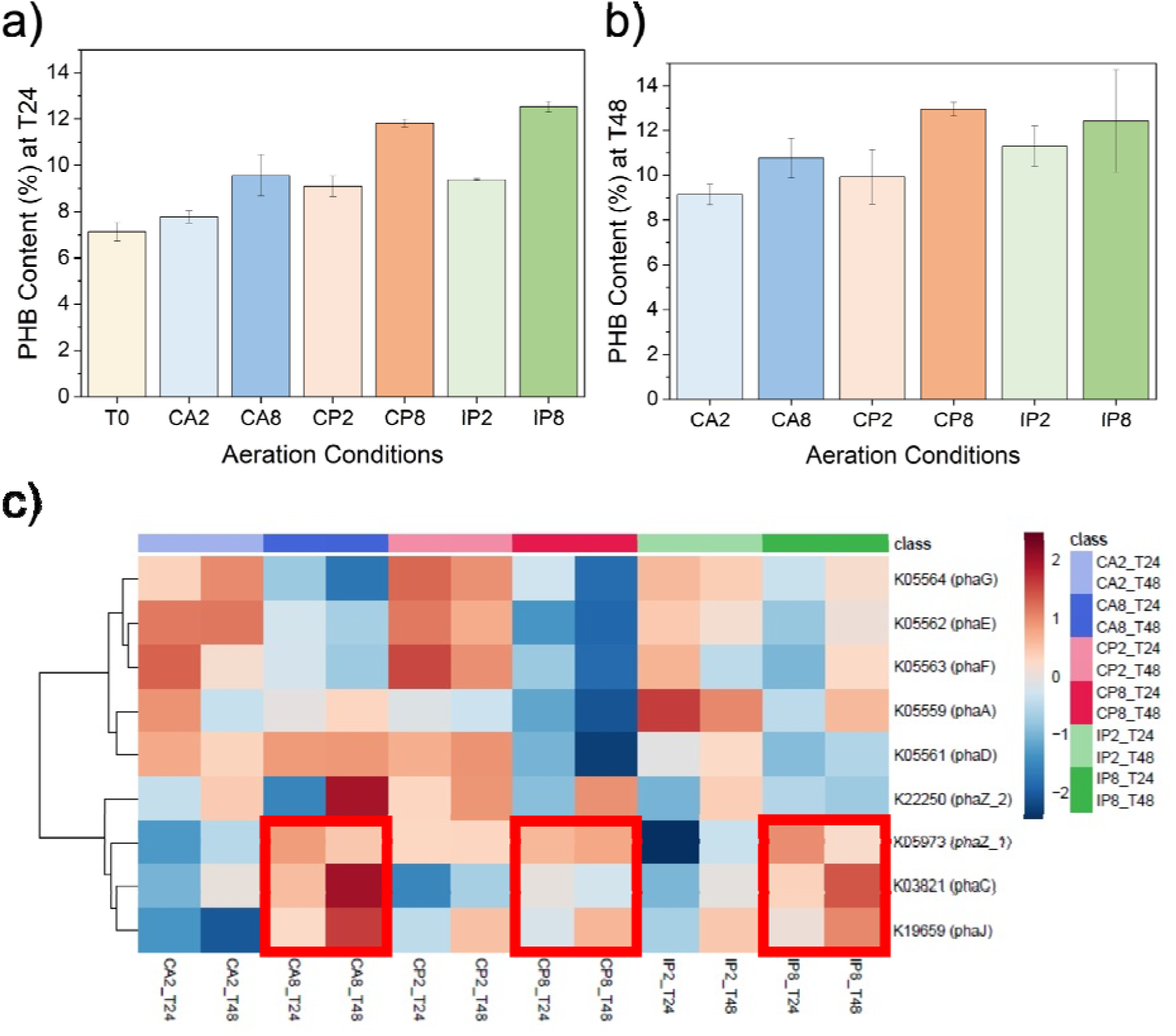
Over-aeration promotes PHB synthesis (derived from fatty acids), utilization and accumulation by microbes: (a, and b) The PHB content in dry biomass samples from activated sludge under six aeration conditions after 24 hours and 48 hours. The error bars represent standard deviations (CA8_T48, CP8_T48, and IP8_T48 have biological duplicates; n=2. Other conditions have biological triplicates; n=3); (c) Abundance of genes associated with the PHA cycle. The data is normalized by Log10 transformation and Pareto scaling. The heatmap show the average value of the same condition. Row clustering according to ‘Ward’ (CA8_T48, CP8_T48, and IP8_T48 have no replicates; n=1. Other conditions have biological triplicates; n=3).

#### 3.4.2 The Genes in PHA Cycle

The synthesis and degradation of PHA involve a series of enzyme-encoding genes. There are three major metabolic pathways for PHA biosynthesis: the acetyl-CoA substrate pathway involving phaA and phaB, encoding β-ketothiolase and acetoacetyl coenzyme A reductase, respectively; the fatty acid β oxidation pathway based on fatty acids involving phaB and phaJ, encoding acetoacetyl CoA reductase and enoyl-CoA hydratase, respectively; and the de novo fatty acid synthesis pathway involving phaG, encoding 3-ketoacyl acyl carrier protein reductase.^53^ All these pathways require phaC-encoded PHA synthase, which polymerizes PHA monomers into PHA polymers in the final step of PHA biosynthesis.^54,55^

In relation to PHA degradation, the gene phaZ plays a critical role as it encodes the PHA depolymerase.^56,57^ Within the activated sludge samples, we detected two variants of the phaZ gene: K05973 (phaZ_1) and K22249 (phaZ_2). Each variant encodes for a different depolymerase, either poly(3-hydroxybutyrate) depolymerase or poly(3-hydroxyoctanoate) depolymerase, responsible for catalyzing the degradation of PHB and poly(3-hydroxyoctanoate) (PHO), respectively.

Based on Figure 6c, in the activated sludge samples, we observed that the genes relevant to the PHA cycle, namely phaJ, phaC, and phaZ, were clustered together, demonstrating a consistent trend of higher abundance under elevated DO conditions. Conversely, other genes, including phaA and phaG, formed a separate cluster.

Combining this with the measured abundance of fatty acids and PHB in the activated sludge samples, we surmise that under the high oxygen levels of the three different aeration patterns, the activated sludge system accumulated more fatty acids intracellularly, which led to a higher abundance of key genes on the pathway producing PHB from fatty acids and a similar increase in genes degrading PHB. This accumulation could be due to the promotion of microbial proficiency in the use of fatty acids for PHB metabolism under high oxygen conditions.

## 4. DISCUSSION

### 4.1 Oxygen Perturbations Promote Protein Release

In this study, the activated sludge microorganisms produced soluble organic matter during metabolism, including extracellular protein, fatty acids, and so forth. There is an intricate interplay between the release of these substances and their accumulation within microbial cells. When the intracellular accumulation of organic matter surpasses the requirements for microbial growth and metabolism, microbes might release some to balance the relationship between intracellular substance accumulation and demand, preventing metabolic and growth restrictions caused by excessive accumulation.^58,59^ Therefore, the released metabolites are then able to represent, to some extent, the metabolic profile of microbes in different environments.

Under different environments, microbes selectively control the utilization efficiency of various amino acids to synthesize needed proteins to adjust cell functions. This is reflected not only in the abundance trend of different types of aminoacyl-tRNA synthetases in IP compared to CA and CP but also in the difference of released protein-like components in SMP. This could be attributed to a range of factors, encompassing metabolic adaptation, ecological competition, and stress response mechanisms. In the face of intermittent oxygen disruptions, microbes might necessitate a higher number of anaerobic metabolic enzymes to navigate the low-oxygen conditions. At the same time, microbes may resort to producing stress proteins to buffer against environmental shifts, aiding their self-preservation and resilience to environmental stressors.

Furthermore, shifts in the microbial community structure might accelerate the phenomenon of differences in the released protein components. Different types of microbes might have distinct metabolic characteristics and protein synthesis capabilities, so community changes could lead to certain types of microbes being more active under specific environmental conditions, thereby affecting the system’s protein synthesis and release.^60,61^

### 4.2 Over-aeration Facilitates Release of Protein and Fatty Acid

Aeration, by supplying oxygen, supports the aerobic respiration processes within the activated sludge system, thereby promoting efficient degradation of organic waste and energy generation by microorganisms. The elevation of environmental oxygen levels in all aeration patterns potentially expedites aerobic metabolic pathways, consequently promoting ATP production that fuels the synthesis of amino acids and fatty acids.^62,63^ In conditions of oxygen scarcity, however, microbes might resort to other metabolic pathways, such as anaerobic glycolysis and respiration. As these pathways generally produce less ATP compared to aerobic pathways, the synthesis rates of amino acids and fatty acids may be comparatively reduced.^64,65^

Under conditions of excessive aeration, compared to the upper limit DO concentration of 2 mg/L, we observed a noticeable augmentation in the intracellular accumulation of amino acids and fatty acids, alongside an increased excretion of protein-like and humic-like substances. On the one hand, a higher oxygen supply provides more oxygen to the microorganisms in the activated sludge, thereby enhancing the synthesis efficiency of amino acids and fatty acids. On the other hand, under conditions of more abundant oxygen, microbes might preferentially exploit simple carbon resource catalyzing for energy generation, leading to a decline in the metabolic utilization efficiency of amino acids and fatty acids.^66^ Furthermore, the intracellular accumulation of fatty acids under these conditions might stimulate PHA synthesis, thereby augmenting the dominance of the fatty acid derived PHA synthesis pathway. Acetyl-CoA, another potential precursor, however, is primarily involved in the energy-producing catabolic pathway rather than PHA synthesis.^67^

Microbial metabolic activity is heightened under excessive aeration, increasing the demand for energy and nutrients. Some SMP excreted by certain microbes can be utilized by others as a nutrient source, thereby fostering material circulation and energy flow within the entire microbial system.^68,69^ However, it is essential to appropriately control oxygen supply during the aeration process. Excessive aeration rates might result in an oxygen surplus, leading to energy waste. Interestingly, promoting the release of SMP from activated sludge can also be achieved by adopting a high oxygen perturbation range strategy. This strategy is an energy-efficient operational approach that does not require a constant oxygen supply, thus averting energy wastage. Additionally, based on oxygen perturbation, continuous and intermittent perturbations can be chosen to regulate wastewater treatment performance and SMP composition, demonstrating the flexibility of this aeration optimization strategy.

## ASSOCIATED CONTENT

### Supporting Information

Detailed materials and methods, the performance of biosystems, 3D-EEM spectral characteristics, the metabolite profiling, the gene abundances, the enzyme abundances (PDF) are described in the supporting information.

### Funding Sources

This study was supported by the Marsden Fund Council from Government funding, managed by Royal Society Te Apārang. [grant number MFP-UOA2018].

## ACKNOWLEDGMENT

We thank Nikki Freed for the assistance with metagenomic sequencing, Martin Middleditch and George Guo for their assistance with metaproteomics analysis, and Saras Green and Alastair Harris for their help with the metabolomics analysis. We also thank Watercare Services Limited for providing the activated sludge culture from the Mangere Wastewater Treatment Plant. The authors acknowledge the use of New Zealand eScience Infrastructure (NeSI) high performance computing facilities, consulting support and/or training services as part of this research. The authors also acknowledge the Centre for eResearch at the University of Auckland for their help in facilitating this research.

## Notes

### Competing Interest Statement

The authors have declared no competing interest.

