## Supplementary for "Assessing Oxygen Perturbation and Over-Aeration Impacts on Soluble Microbial Products (SMP) Production and Release in Activated Sludge Systems"

### for

*Xueyang Zhou[1,2], Bharat Manna[1,2], Boyu Lyu[1,2], and Naresh Singhal[1,2]* *
[1]Department of Civil and Environmental Engineering, University of Auckland, Auckland 1010, New Zealand
[2]Water Research Centre, University of Auckland, Auckland 1010, New Zealand

#### S1. SUPPLEMENTARY METHODS

##### S1.1 Experimental setup and operation

**Table S1.** Composition of trace element solution

| Chemical | Concentration (g/L) |
|---|---|
| EDTA | 2.50 |
| $ZnSO_4 \cdot 7H_2O$ | 1.10 |
| $CoCl_2 \cdot 6H_2O$ | 0.800 |
| $MnCl_2 \cdot 4H_2O$ | 2.55 |
| $MgSO_4 \cdot 7H_2O$ | 20.0 |
| $CuSO_4 \cdot 5H_2O$ | 0.860 |
| $(NH_4)_6Mo_7O_{24} \cdot 4H_2O$ | 0.0680 |
| $CaCl_2 \cdot 2H_2O$ | 2.75 |
| $FeSO_4 \cdot 7H_2O$ | 2.57 |

**Table S2.** Composition of the concentrated artificial wastewater

| Chemical | Concentration |
|---|---|
| Methanol (mL/L) | 26.936 |
| $NH_4Cl$ (g/L) | 24.457 (IP); 30.571 (CP or CA) |
| $KH_2PO_4$ (g/L) | 2.7550 |
| $K_2HPO_4$ (g/L) | 2.7550 |
| $NaHCO_3$ (g/L) | 76.800 |

#### S2. Results

##### S2.1. The biosystem performance under different aeration conditions

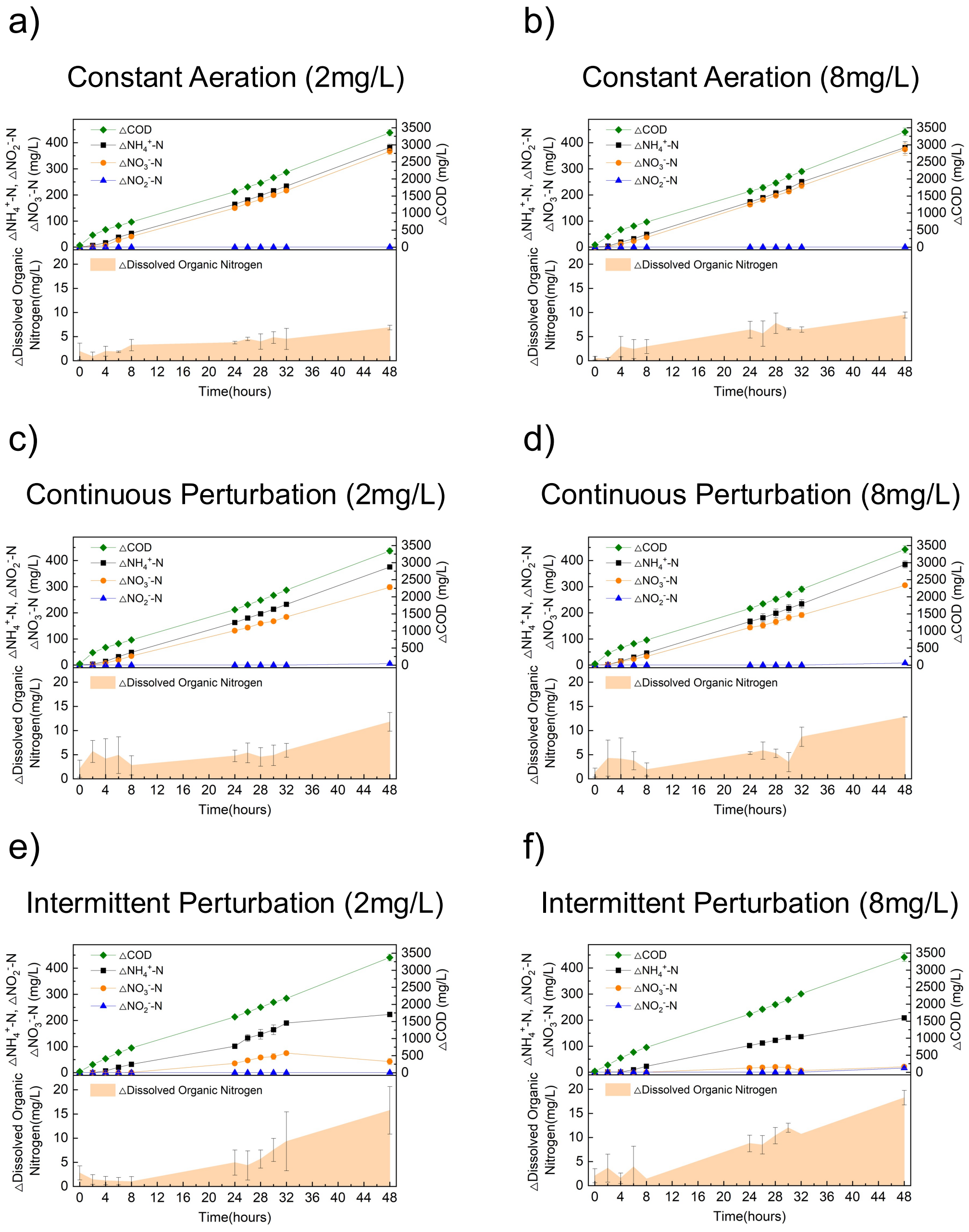


**Figure S1.** The biosystem performance and dissolved organic nitrogen accumulation in activated sludge systems exposed to different aeration patterns. Error bars represent standard deviations (CA2, CP2, and IP2 have biological triplicates; n=3. CA8, CP8, and IP8 have biological duplicates; n=2)

**Table S3.** Carbon and nitrogen conversion efficiency for 48 hours running under different aeration patterns

| | CA2 | CA8 | CP2 | CP8 | IP2 | IP8 |
|---|---|---|---|---|---|---|
| COD removal rates (mg/L/hour) | 63.63 | 64.42 | 63.62 | 64.85 | 65.87 | 67.64 |
| $NH_4^+$-N removal rates (mg/L/hour) | 7.78 | 8.02 | 7.76 | 7.94 | 5.25 | 4.64 |
| $NO_2^-$-N accumulation rates (mg/L/hour) | 0.01 | 0.01 | 0.06 | 0.09 | 0.01 | 0.19 |
| $NO_3^-$-N accumulation rates (mg/L/hour) | 7.51 | 7.90 | 6.25 | 6.46 | 1.60 | 0.53 |

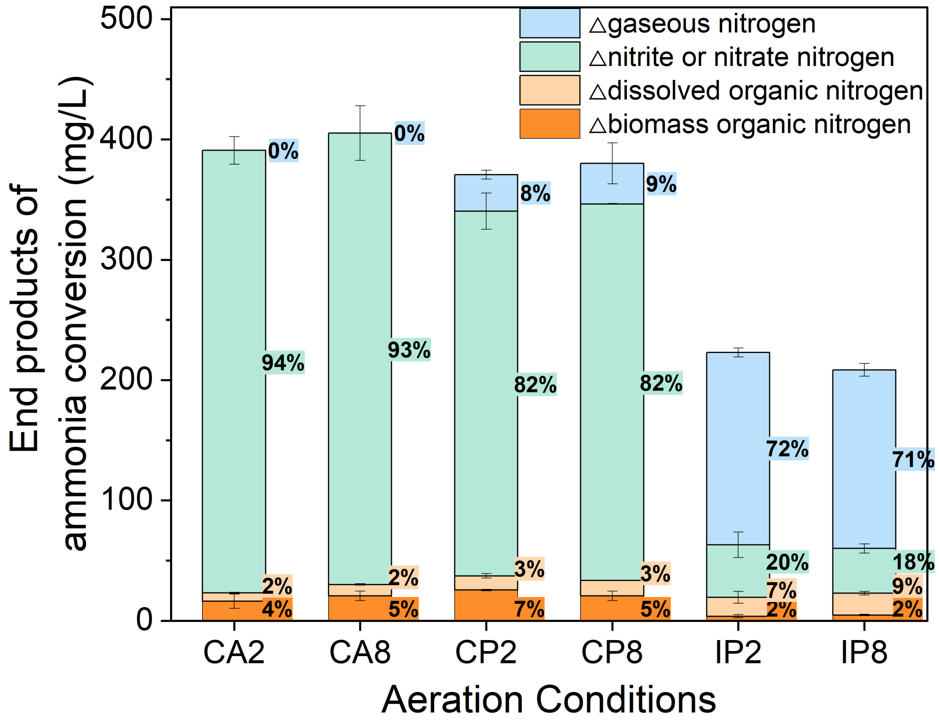


**Figure S2.** The amount of different end products converted from ammonia by the activated sludge system exposed to different aeration strategies for 48 hours. Error bars

represent standard deviations (CA2, CP2, and IP2 have biological triplicates; n=3. CA8, CP8, and IP8 have biological duplicates; n=2)

##### S2.2. The dissolved organic matters measurement by 3D-EEM

**Table S4.** The properties of 4 components identified by Openfluor

| Component | Excitation maximum | Emission maximum | Assignment | References |
|---|---|---|---|---|
| C5 | 351 nm | 441 nm | Humic-like compound | (Shutova et al., 2014) |
| C6 | 285 nm | 339 nm | Protein-like compound | (Lapierre & del Giorgio, 2014) |
| C7 | 282 nm | 336 nm | Protein-like compound | (Dalmagro et al., 2019) |
| C8 | 278 nm | 524 nm | Fulvic compound | (Hong et al., 2021) |

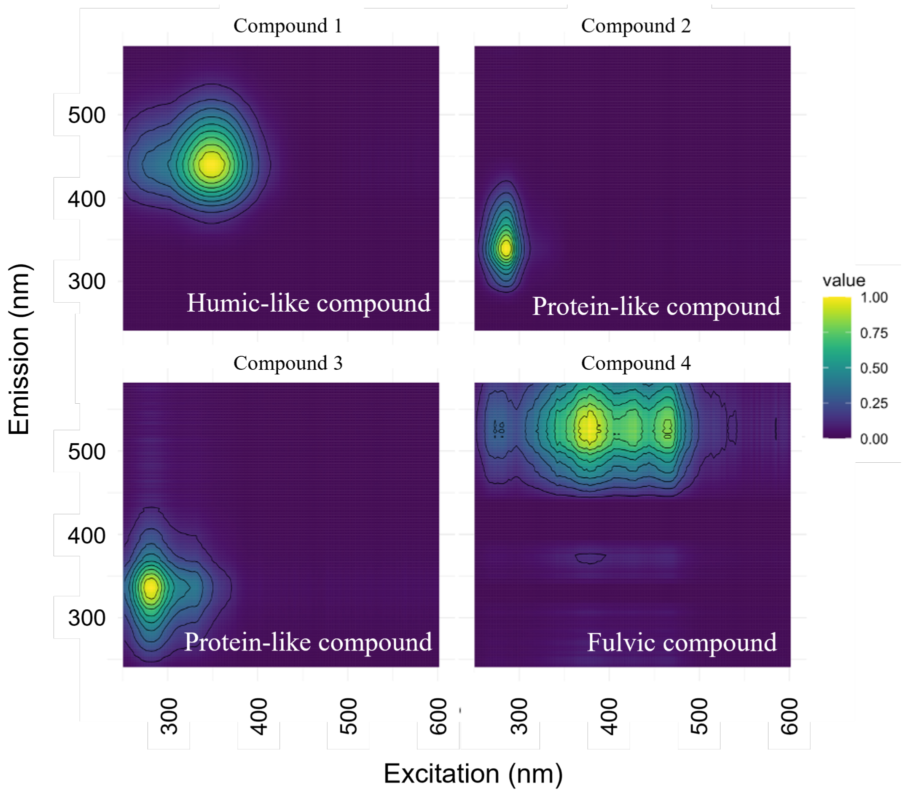


**Figure S3.** fDOM component spectral characteristics in extracellular components were identified by the PARAFAC model

#### S2.3. The amino acid measurement by MCF derivatization metabolomics

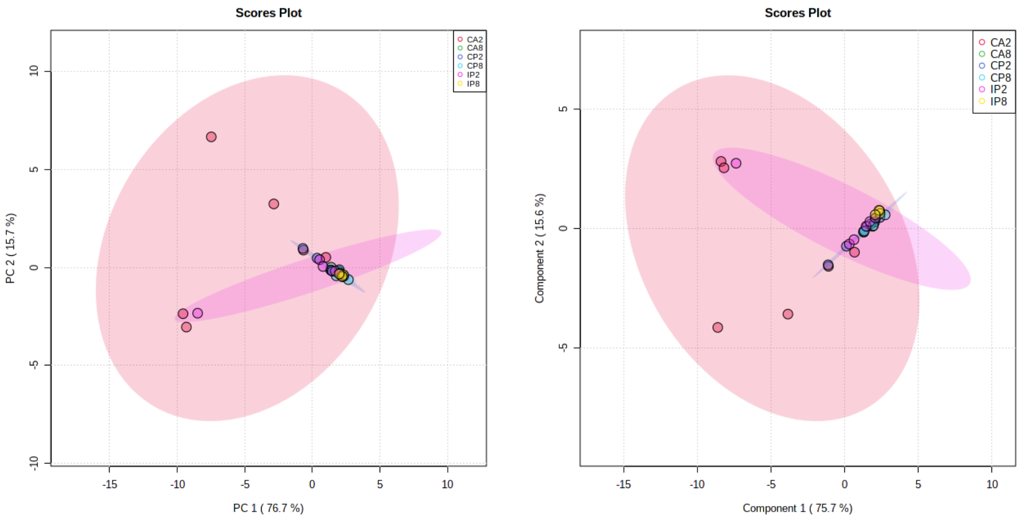


**Figure S4.** PCA and PLS-DA score plot of intracellular amino acids related metabolic profiling

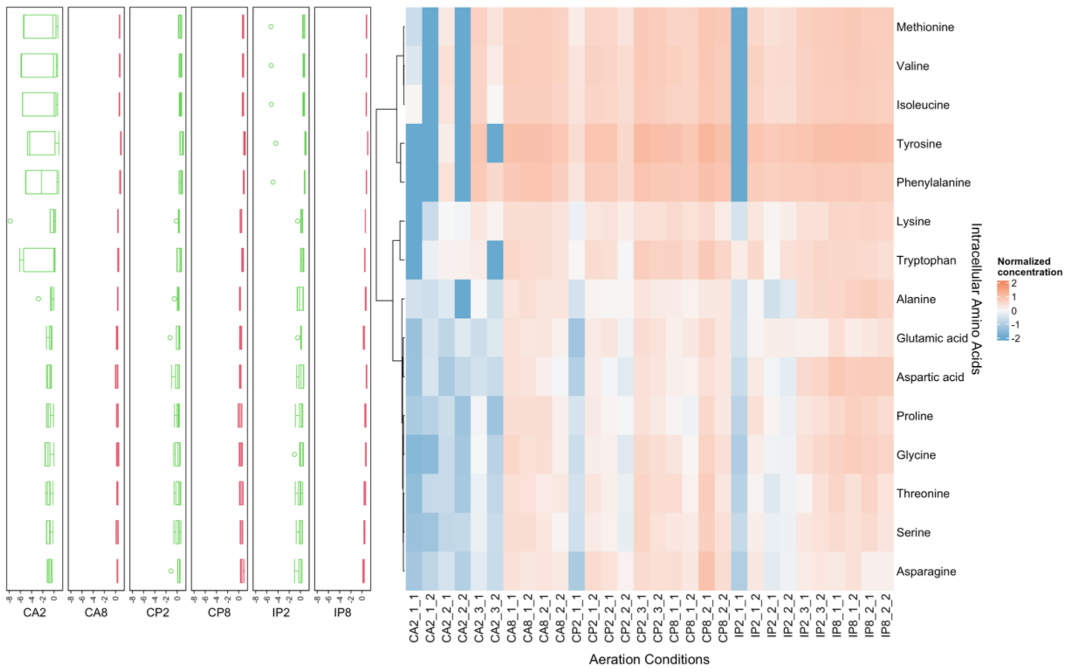


**Figure S5.** The boxplot and heatmap of intracellular amino acids related metabolic profiling. The data is normalized by internal standard, Log10 transformation, and Pareto scaling. Row clustering according to 'Ward'. The box chart on the left shows the average value of the same condition (CA2, CP2, and IP2 have biological triplicates; n=3. CA8, CP8, and IP8 have biological duplicates; n=2)

##### S2.4. The abundance of tRNA synthetases by metaproteomics

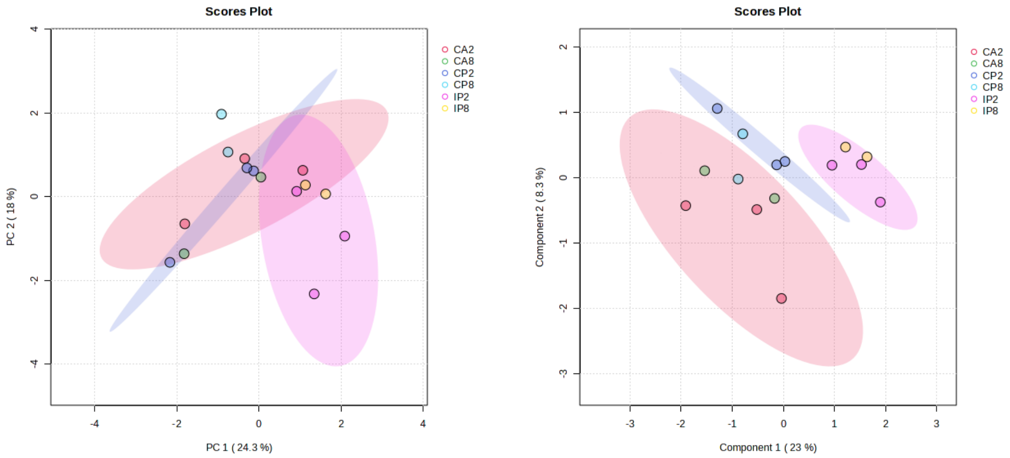


**Figure S6.** PCA and PLS-DA score plot of tRNA synthetases in different metaproteomic samples

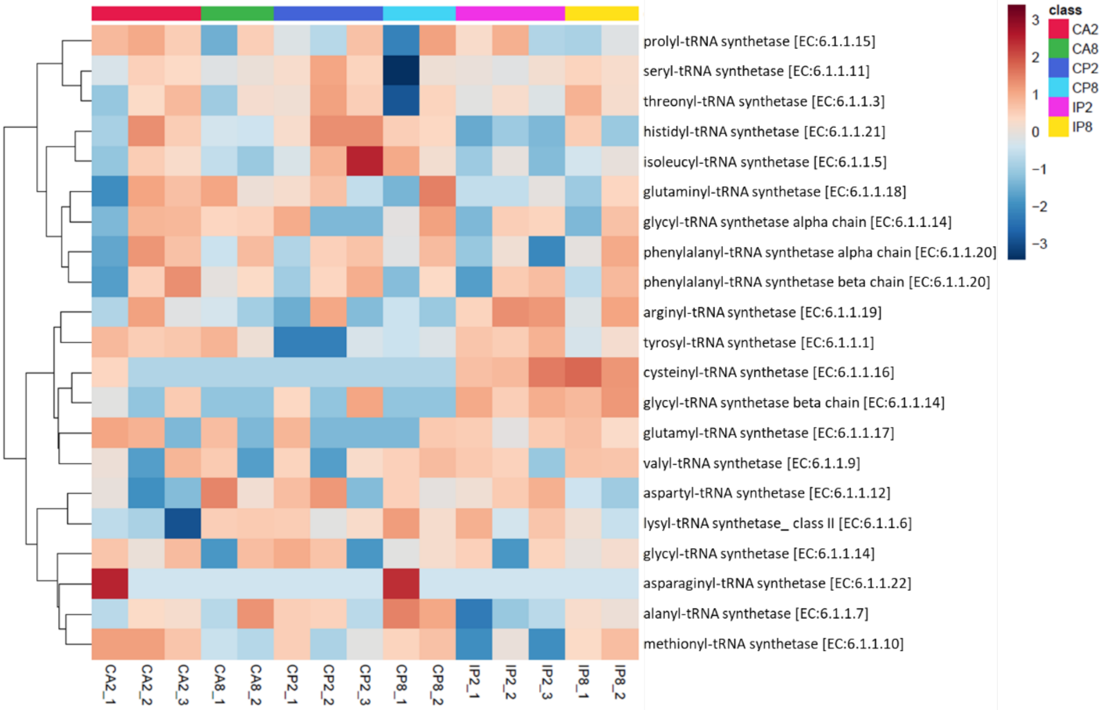


**Figure S7.** The heatmap of tRNA synthetases in different metaproteomic samples.

The data is normalized by Log10 transformation, and Pareto scaling. Row clustering according to 'Ward' (CA2, CP2, and IP2 have biological triplicates; n=3. CA8, CP8, and IP8 have biological duplicates; n=2)

##### S2.5. The fatty acid measurement by FAMEs derivatization metabolomics

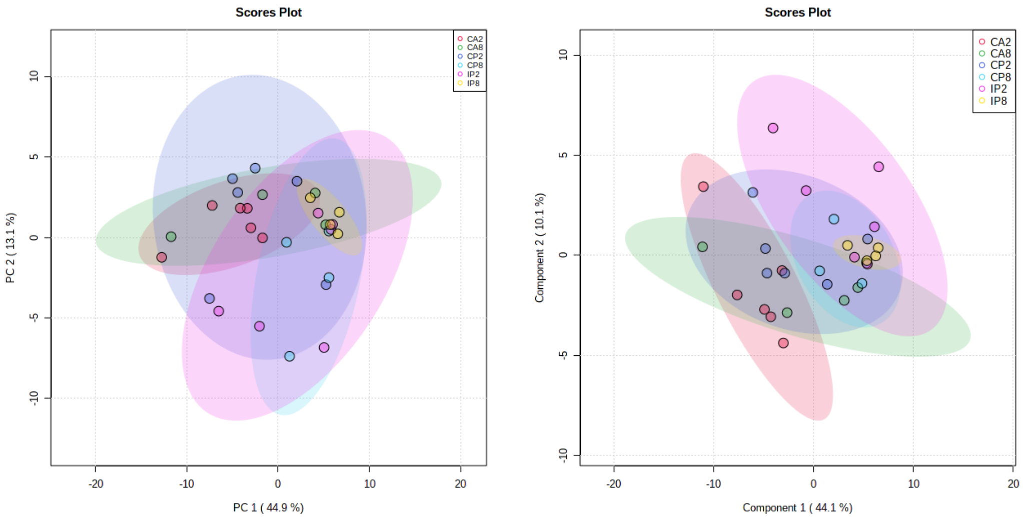


**Figure S8.** PCA and PLS-DA score plot of intracellular fatty acids related metabolic profiling

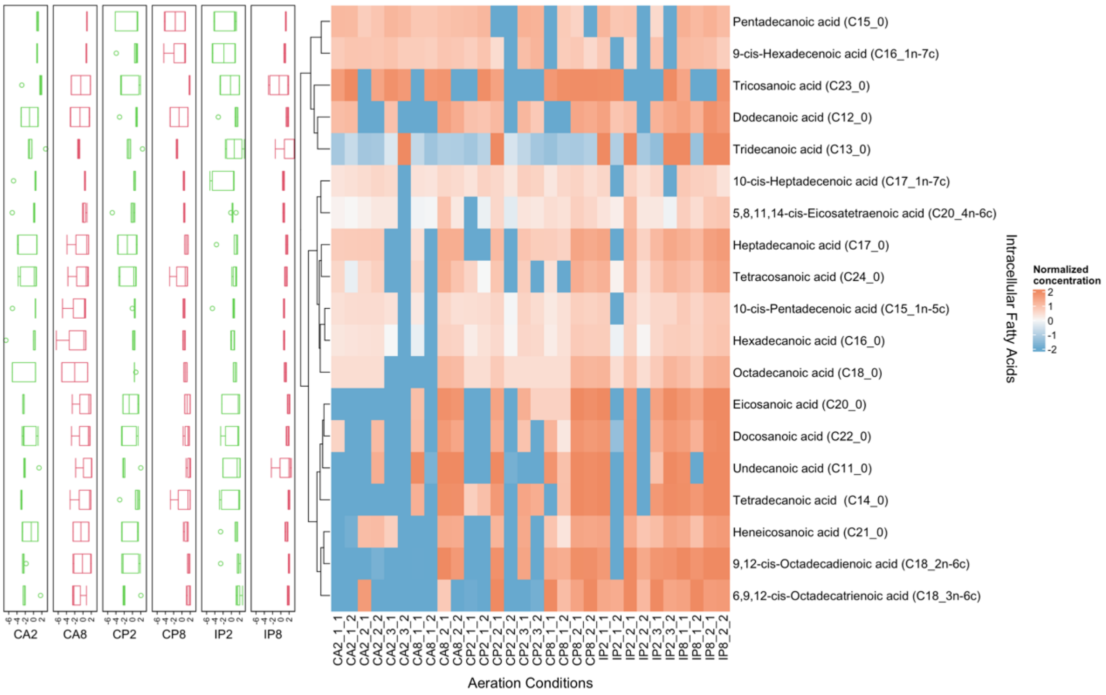

**Figure S9.** The boxplot and heatmap of intracellular fatty acids related metabolic profiling. The data is normalized by internal standard, Log10 transformation, and Pareto scaling. Row clustering according to 'Ward'. The box chart on the left shows the average value of the same condition (CA2, CP2, and IP2 have biological triplicates; n=3. CA8, CP8, and IP8 have biological duplicates; n=2)

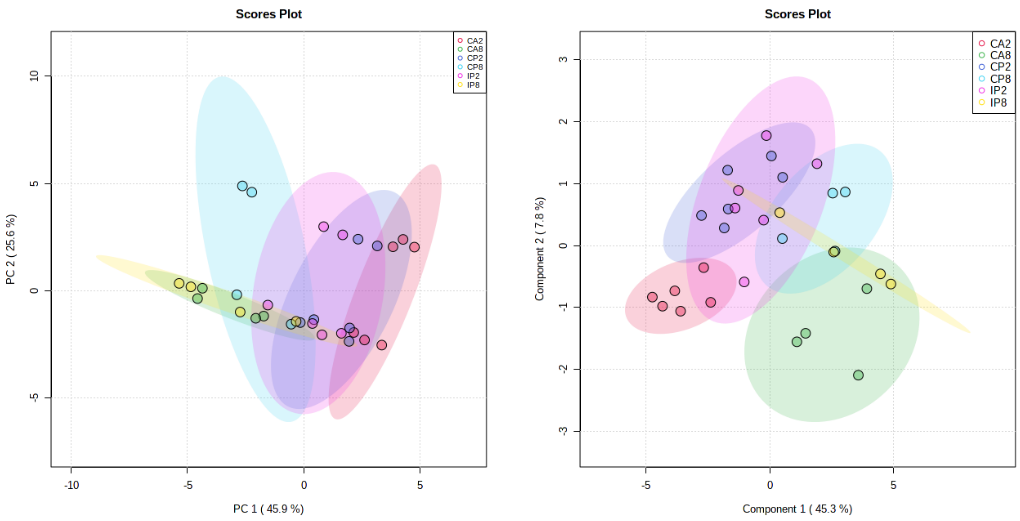


**Figure S10.** PCA and PLS-DA score plot of extracellular fatty acids related metabolic profiling

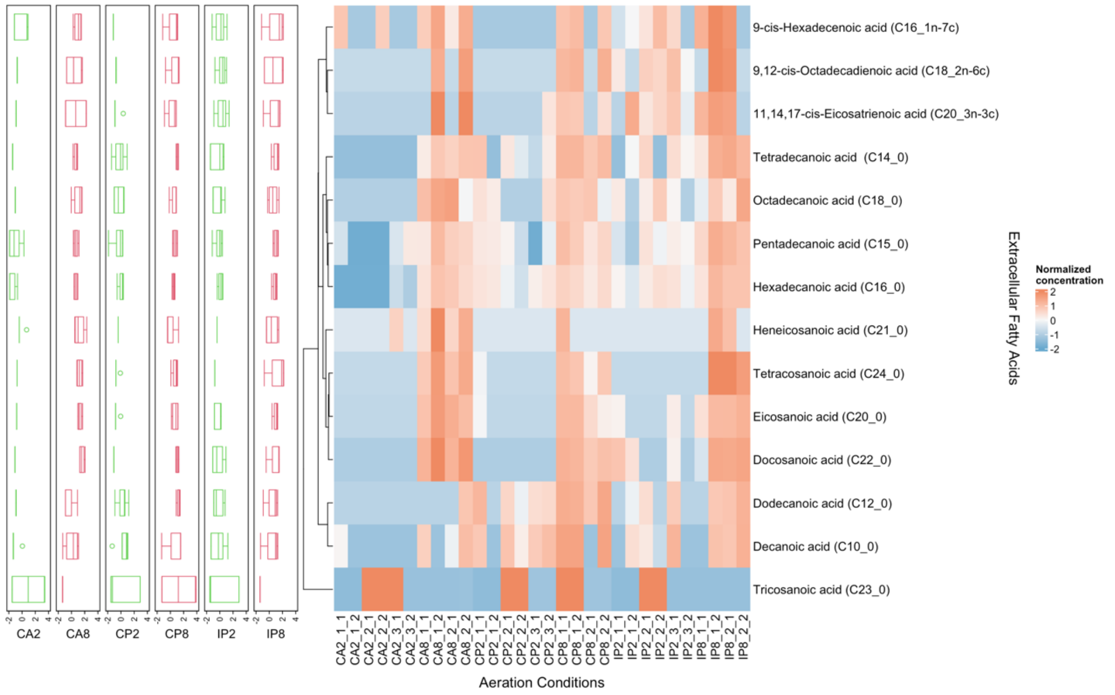


**Figure S11.** The boxplot and heatmap of extracellular fatty acids related metabolic profiling. The data is normalized by internal standard, Log10 transformation, and Pareto scaling. Row clustering according to 'Ward'. The box chart on the left shows the average value of the same condition (CA2, CP2, and IP2 have biological triplicates; n=3. CA8, CP8, and IP8 have biological duplicates; n=2)

##### S2.6. The genes related with fatty acids biosynthesis metabolism

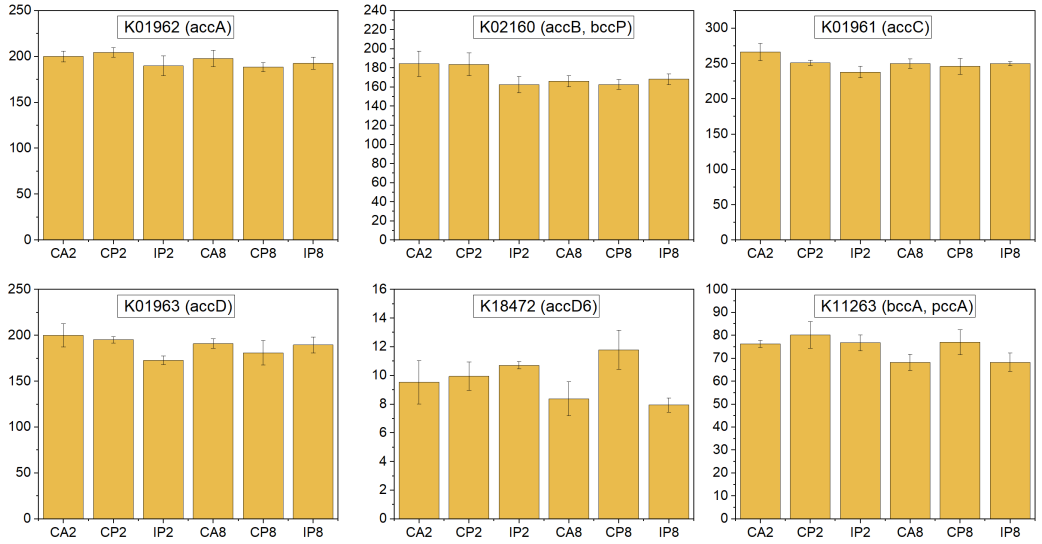


**Figure S12.** The transcripts per million (TPM) of the genes related to acetyl-CoA carboxylase (EC:6.4.1.2) of activated sludge samples taken at 24 hours under different aeration conditions. The error bars represent standard deviations (biological triplicates; n=3)

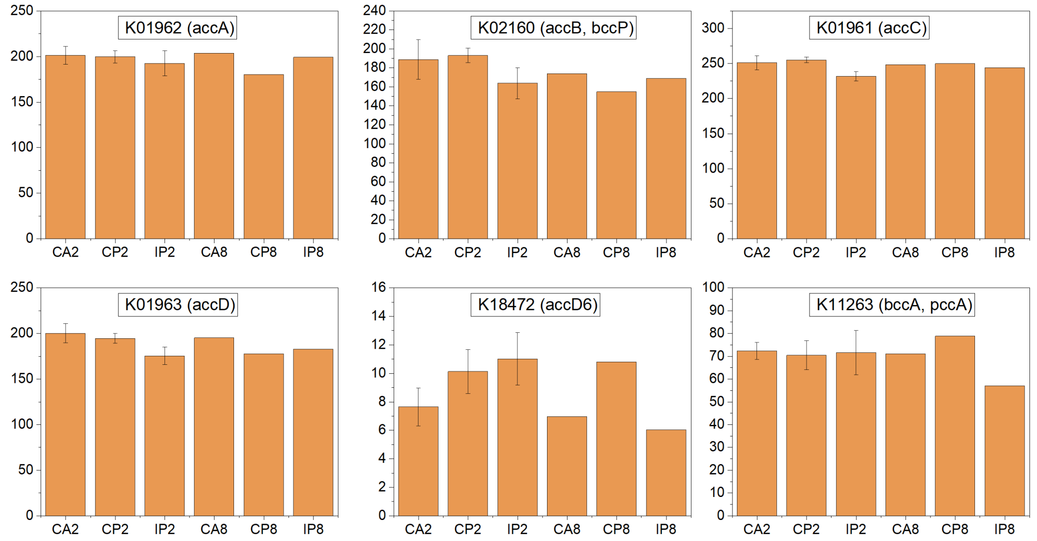


**Figure S13.** The transcripts per million (TPM) of the genes related to acetyl-CoA carboxylase (EC:6.4.1.2) of activated sludge samples taken at 48 hours under different aeration conditions. The error bars represent standard deviations (CA2, CP2, and IP2 have biological triplicates; n=3. CA8, CP8, and IP8 have no replicates; n=1)

#### S2.7. The genes related with fatty acids utilization metabolism

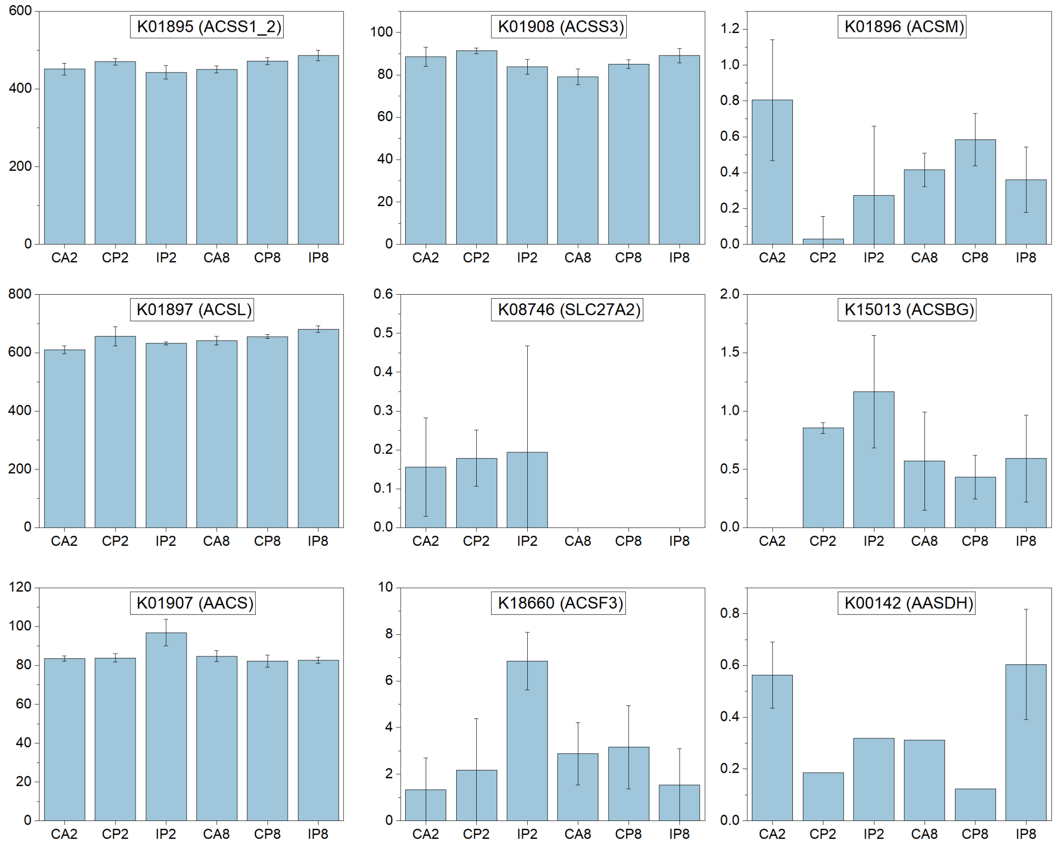


**Figure S14.** The transcripts per million (TPM) of the genes related to acyl-CoA synthetase family members of activated sludge samples taken at 24 hours under different aeration conditions. The error bars represent standard deviations (biological triplicates; n=3)

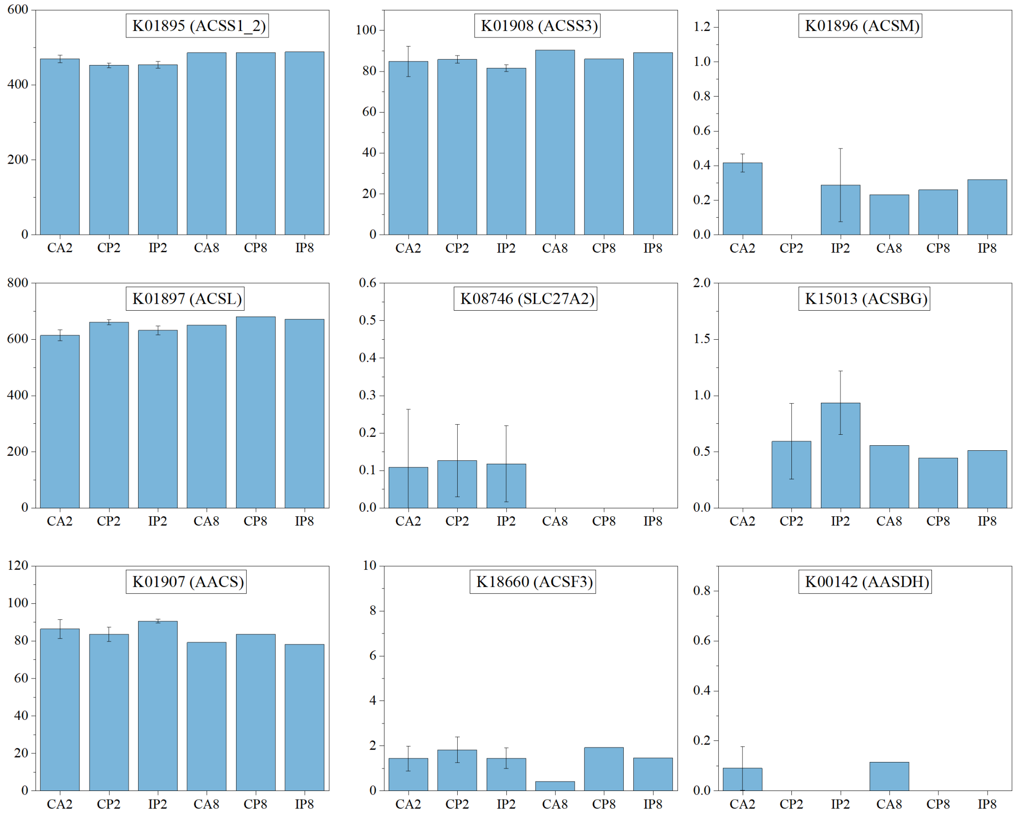


**Figure S15.** The transcripts per million (TPM) of the genes related to acyl-CoA synthetase family members of activated sludge samples taken at 48 hours under different aeration conditions. The error bars represent standard deviations (CA2, CP2, and IP2 have biological triplicates; n=3. CA8, CP8, and IP8 have no replicates; n=1)

**Table S5.** Acyl-CoA synthetase identified in activated sludge samples

| ACS subfamily | Gene ID | Official symbol | Official name |
|---|---|---|---|
| Short-chain | K01895 | ACSS1 | Acyl-CoA synthetase short-chain family member 1 |
| | K01908 | ACSS3 | Acyl-CoA synthetase short-chain family member 3 |
| Medium-chain | K01896 | ACSM | Acyl-CoA synthetase medium-chain family member |
| Long-chain | K01897 | ACSL | Acyl-CoA synthetase long-chain family member |
| Very long-chain | K08746 | SLC27A2 | Solute carrier family 27 (fatty acid transporter), member 2 |
| 'Bubblegum' | K15013 | ACSBG | Acyl-CoA synthetase bubblegum family member |
| Other | K01907 | AACS | Acetoacetyl-CoA synthetase |
| | K18660 | ACSF3 | Acyl-CoA synthetase family member 3 |
| | K00142 | AASDH | Aminoadipate-semialdehyde dehydrogenase |

#### S2.8. The detoxification of very long-chain fatty acids

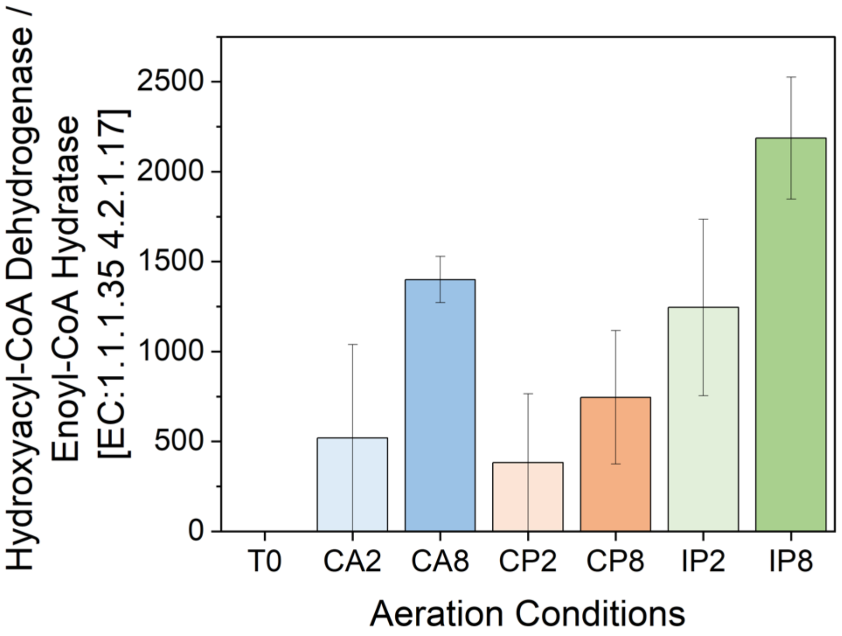


**Figure S16.** The enzyme abundance of the bifunctional protein with both enoyl-CoA hydratase and hydroxyacyl-CoA dehydrogenase under different aeration conditions. The error bars represent standard deviations (CA2, CP2, and IP2 have biological triplicates; n=3. CA8, CP8, and IP8 have biological duplicates; n=2)

#### S2.9. The genes of PHA cycle

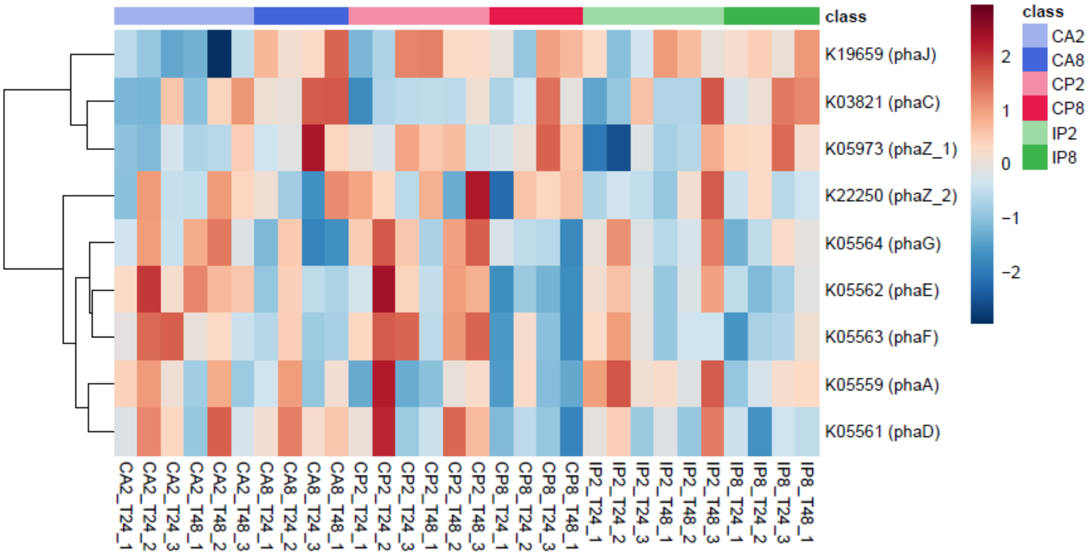


**Figure S17.** The abundance of genes related to PHA cycle. The data is normalized by Log10 transformation and Pareto scaling. Row clustering according to 'Ward'
